# Cognitive and brain function enhancement in Gen X group after personalized, AI supervised EEG-neurofeedback training

**DOI:** 10.1101/2025.11.17.688986

**Authors:** Jacek Rogala, Urszula Malinowska, Michał Ociepka, Jakub Wojciechowski, Joanna Zych, Przemysław Tryc, Anna Kołodziejak, Paweł Ogniewski, Paweł Niedbalski, Jan Skorupski, Adam Chuderski

## Abstract

**Background:** Interventions supporting medical care and enhancing quality of life in neurodegenerative or age-related cognitive decline are strongly needed. Electroencephalographic (EEG) neurofeedback can enable users to modulate their brain activity through real-time feedback. However, evidence for its clinical effectiveness remains inconclusive, partly due to limited personalization and insufficient task relevance in existing protocols.

**Objective:** We tested whether personalized EEG neurofeedback supervised by deep neural networks (DNNs) can enhance cognitive performance in older adults.

**Methods:** Fifty-seven healthy adults aged 41–64 (31 women), including a sham-feedback control group, completed a personalized neurofeedback protocol with DNNs fine-tuned to individual EEG patterns. The procedure included pre- and post-training assessments using a transitive reasoning task, three diagnostic sessions to adapt the DNN to each participant, and 10–11 neurofeedback sessions based on a gamified delayed-match-to-sample paradigm.

**Results:** The training group showed robust gains across all three variants of the reasoning task (each p < .01), whereas the sham group improved only on the easiest variant. Groups did not differ at pretest; however, at posttest the training group outperformed the sham group on all task conditions (each p < .03), showing also a larger neural effort (lower alpha band power) and increased beta and gamma band connectivity (higher phase lag index).

**Conclusion:** Personalized, task-oriented neurofeedback guided by individually fine-tuned DNNs can produce cognitive enhancement after relatively few sessions. The proposed Task-Pretrained, Subject-Finetuned Neurofeedback (TPSF-NF) framework is scalable to other cognitive domains in future research.

## Introduction

The global rise in neurodegenerative diseases is linked to both the aging of the population and the increasing levels of stress. According to The Lancet Neurology [1], neurological disorders were responsible for 10.2% of all years of life lost in 2015, ranking as the leading cause of such loss and the second leading cause of death. This burden is especially acute in individuals aged 55 to 74 and is expected to escalate as populations continue to age and neurological conditions become more prevalent across all age groups. Consequently, addressing the treatment, rehabilitation, and prevention of nervous system disorders has emerged as a critical health priority.

In this context, there is a growing need for therapeutic strategies that not only complement traditional medical treatments but also enhance the quality of life for individuals with neurodegenerative conditions or those seeking to preserve or improve cognitive functions. One such approach is electroencephalographic (EEG) neurofeedback—a technique that enables individuals to regulate their brain activity through real-time feedback. Despite its widespread use, the clinical effectiveness of EEG neurofeedback remains inconclusive. Emerging evidence suggests that this may stem, in part, from the application of standardized protocols to populations characterized by high variability, particularly older adults experiencing cognitive decline ([2], [3], [4]).

Crucially, many of these neurofeedback protocols are developed and validated using data from young adults, whose brain activity, functional connectivity, and cognitive performance differ substantially from those of older individuals ([5], [6], [7], [8], [9], [10], [11], [12], [13], [14]). As a result, protocols based on generalized associations between EEG activity and cognitive functions may not be effective when applied to individuals with distinct neurophysiological and cognitive profiles.

To improve therapeutic outcomes, EEG neurofeedback should be personalized— grounded in the individual’s specific EEG patterns recorded during tasks related to the targeted cognitive function (for review see [15]). However, the vast number of EEG features and their complex, individual-specific associations with cognitive processes make this task difficult using conventional analytical methods.

In recent years, advances in algorithms, data availability, and computing power have driven the rapid growth of machine learning, particularly deep neural networks (DNNs). These models offer promise for addressing the limitations of traditional EEG analysis. DNNs can automatically extract a broad array of features and optimize feature selection through end-to-end learning, enhancing EEG classification performance ([16], [17]). Moreover, unlike traditional statistical methods, DNNs analyze all features concurrently, enabling them to detect complex patterns in EEG activity associated with specific neurophysiological or cognitive states ([18]). This makes them well-suited to guide personalized neurofeedback interventions.

In the present study, we used DNNs to supervise and tailor EEG neurofeedback training to individual participants. The employed machine learning model was designed specifically to identify EEG features associated with individual memory processes for neurofeedback application [18]. Current study provides evidence that personalized, task-oriented neurofeedback training, guided by a tailored DNN, can drive cognitive improvements after a limited number of training sessions. The proposed concept of Task-Pretrained, Subject-Finetuned Neurofeedback (TPSF-NF) can be easily adapted to improvement of various cognitive functions, given access to task specific EEG data. Having in mind our aim of enhancing cognitive performance in older adults—those who stand to benefit most from such interventions—this study examined adults aged 41 to 64, spanning approximately the range of Generation X (i.e., individuals born in the 60s and 70s). Our main goal was to test whether the TPSF-NF protocol could lead to cognitive enhancement across three working memory tasks varying in complexity (maintenance, updating, and relation processing), as compared to a sham control group. To foretell the results, TPSF-NF effectively enhanced cognitive processing in complex/difficult, but not in simpler/easier, working memory task conditions.

## Methods

### Participants

To determine the required sample size, a power analysis was conducted using *G*Power* [19] for a repeated-measures within–between interaction ANOVA with two measurement points, an alpha level of 0.05, and a medium effect size (*f* = 0.25). The analysis suggested a minimum total sample size of 54 participants. For this study, 60 participants were initially recruited through announcements at local employment agencies in a large Central European city. Ultimately, 57 healthy adults (31 women) completed the experiment. This sample size was comparable to or larger than those in similar neurofeedback studies ([20]: *n* = 32; [21]: *n* = 60; [22]: *n* = 40; [23]: *n* = 30). The participants’ mean age was 53.0 years (*SD* = 7.1, range: 41-64). All individuals were right-handed and had normal or corrected-to-normal vision. Participants were excluded if they had a history of mental disorders, dementia or related neurodegenerative conditions, substance abuse, and stroke.

Participants were randomly assigned to one of two groups: 28 to the real neurofeedback training group and 29 to the control (sham) group (pretended neurofeedback). Group allocation was based on a coded identification system, developed by researchers who did not take part in later stages of the procedure.

During training, the participant’s code was entered into the proprietary Digitrack system, which automatically selected the appropriate training protocol—real or sham—according to the predefined assignment. In each session, a participant was assigned randomly to one of the four research assistants supervising the procedure, to avoid any influence of a given assistant. A third, independent team of researchers conducted the data analysis. Therefore, personnel responsible for group assignment, training supervision, and data analysis remained separate. This structure guaranteed a fully double-blind experimental design.

All participants received monetary compensation in local currency (amounting to approximately 300 USD) for taking part in up to 18 neurofeedback sessions, including three diagnostic sessions. The study was approved by the local ethics committee. All participants provided written informed consent prior to participation.

### Neurofeedback procedure

The neurofeedback protocol was implemented through a computer game based on a modified delayed match-to-sample (DMTS) paradigm ([18], [24]). The game was developed in the Unity programming environment and integrated within the DigiTrack system (Elmiko BIOSIGNALS, Milanówek, Poland). EEG DigiTrack is a certified medical-grade EEG biofeedback platform that enables flexible design and implementation of EEG-based training protocols. The game was presented on a Windows-based computer equipped with a 24-inch monitor.

In the proper neurofeedback training version, each trial followed the three standard DMTS stages—sample, delay, and choice—embedded within the gameplay. During the *sample* phase, a silhouette of an enemy spaceship (the target) was displayed. After a 5-second *delay* period, the *choice* phase began, in which a spaceship silhouette appeared in front of the player’s cabin (see Fig. 1). Each trial lasted 11 seconds. Participants were asked to determine whether the presented silhouette matched the previously shown target. If it matched, they were instructed to “shoot” the ship; if not, they were to activate a protective force field, identifying it as friendly. For each trial, participants were instructed to reproduce the mental state associated with correct responses during the preceding diagnostic sessions (described below).

**Fig. 1.**
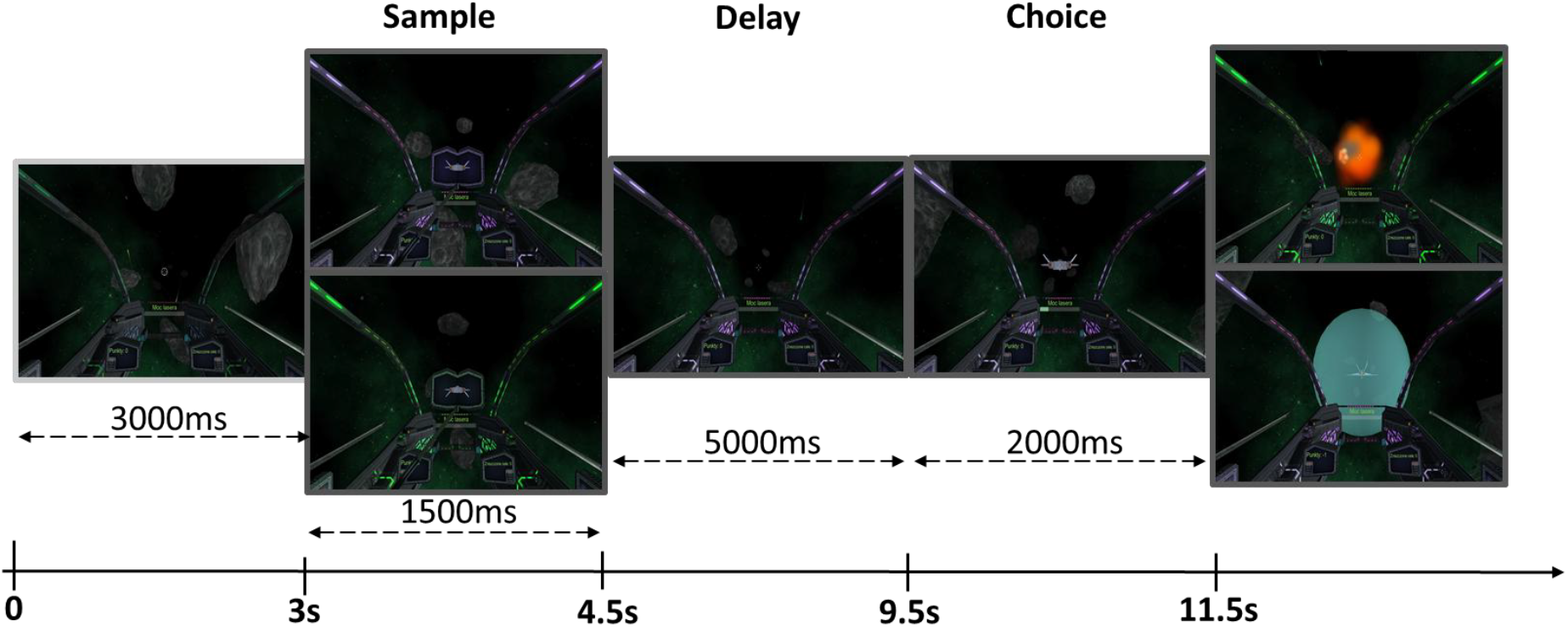
DMTS task scheme. On the top, an example with a matched silhouette (enemy ship to shoot down), and below a version with no matched target (to protect by a force field).

Before the main neurofeedback training, three diagnostic sessions were conducted (on separate days, 2-3 days apart) to decode participants’ EEG activity while performing the game. During the diagnostic phase, participants used a keyboard to play the game while their EEG was recorded. Data from these sessions were used to identify EEG patterns related to memory retention (correct trials only) and to fine-tune previously trained deep neural network (DNN) models. The diagnostic version included two trial types: *difficult* trials—identical to those used in the main training, requiring memory retention—and *easy* trials, which did not involve memory retention. In easy trials, a tilted meteor replaced the enemy spaceship during the choice phase, and participants indicated the meteor’s tilt direction. Each diagnostic session comprised 100 trials (50 of each type) presented in random order. Trial types were visually distinguished by different ambient cabin lighting colors.

Both the neurofeedback and the control group followed exactly the same procedures, except for one control parameter (i.e., the classifier result) which in the control group was distorted by a pseudorandom generator by adding a random noise to the true value, resulting in a 60% chance of success. To prevent undesired reactions of the personnel which could affect participants’ responses, all parameters on the personnel’s screen were always real, consistent with the electrode recordings. This also served to hide the information about the sham from the personnel.

The neurofeedback procedure as well as 10 proper training sessions were carried out with the use of the 19-channel DigiTrack device by Elmiko (using the ExG-32 headbox manufactured by ELMIKO BIOSIGNALS, Milanówek, Poland), with 21 electrodes arranged in a 10-20 system, referenced to the right ear. According to the routine medical procedure the impedance of the electrodes was always kept below 10 KΩ. The sampling frequency was 500 Hz. The training process during the training sessions was supervised by a DNN pre-trained on the EEG recordings acquired during three manual DMTS sessions of 87 participants in our previous study [18], and tuned to each new participant using data from diagnostic sessions.

### Machine Learning based neurofeedback protocol

The DNN employed for supervising neurofeedback training in the current experiment was the Hybrid model, specifically developed to support EEG-based neurofeedback procedures [18]. The Hybrid model was trained in a two-step procedure: general training followed by fine-tuning to specific participants using raw data from diagnostic sessions. For the general training phase, EEG data underwent standard preprocessing to eliminate artifacts that might bias classification and, by extension, the neurofeedback intervention. The initial model training was conducted using the PyTorch library on a Unix-based operating system. The fine-tuning phase was implemented in C++ and executed within the targeted Windows environment. Fine-tuning was performed directly on raw, unprocessed EEG signals to maximize ecological validity and improve applicability during therapeutic sessions.

During general training, the EEG signals were transformed into time-frequency representations using the Morlet wavelet transform (wave number = 7), applied at central frequencies of 3, 5, 8, 11, 15, 20, 25, 30, and 35 Hz. This transformation produced a time-frequency map for each EEG channel. These individual maps were then stacked across channels to construct a three-dimensional tensor with dimensions *(channels × frequency × time)*, representing the model input as E × F × T, where *E* is the number of electrodes, *F* the number of frequency bands, and *T* the number of time steps.

Within each computational block of the DNN, a convolutional layer was followed by a ReLU (Rectified Linear Unit) activation function and batch normalization [25]. The model incorporated two convolutional layers, each comprising 8 kernel channels per location. To capture temporal features at multiple scales, these layers were arranged into three parallel branches, with each branch employing different kernel sizes along the time dimension [25]. The outputs of these branches were aggregated using an average pooling operation applied across the entire trial duration. Finally, the pooled outputs were concatenated, yielding a 24-dimensional feature representation of the input (8 features per branch).

This 24-dimensional representation was subsequently passed through two fully connected hidden layers. The final trained output of the ‘concatenation’ block was then used as a fixed feature extractor. This representation served as the input to individual logistic regression models, which were trained on participant-specific data obtained across three diagnostic sessions. The whole Hybrid model architecture is presented in Fig. 2.

**Fig. 2.**
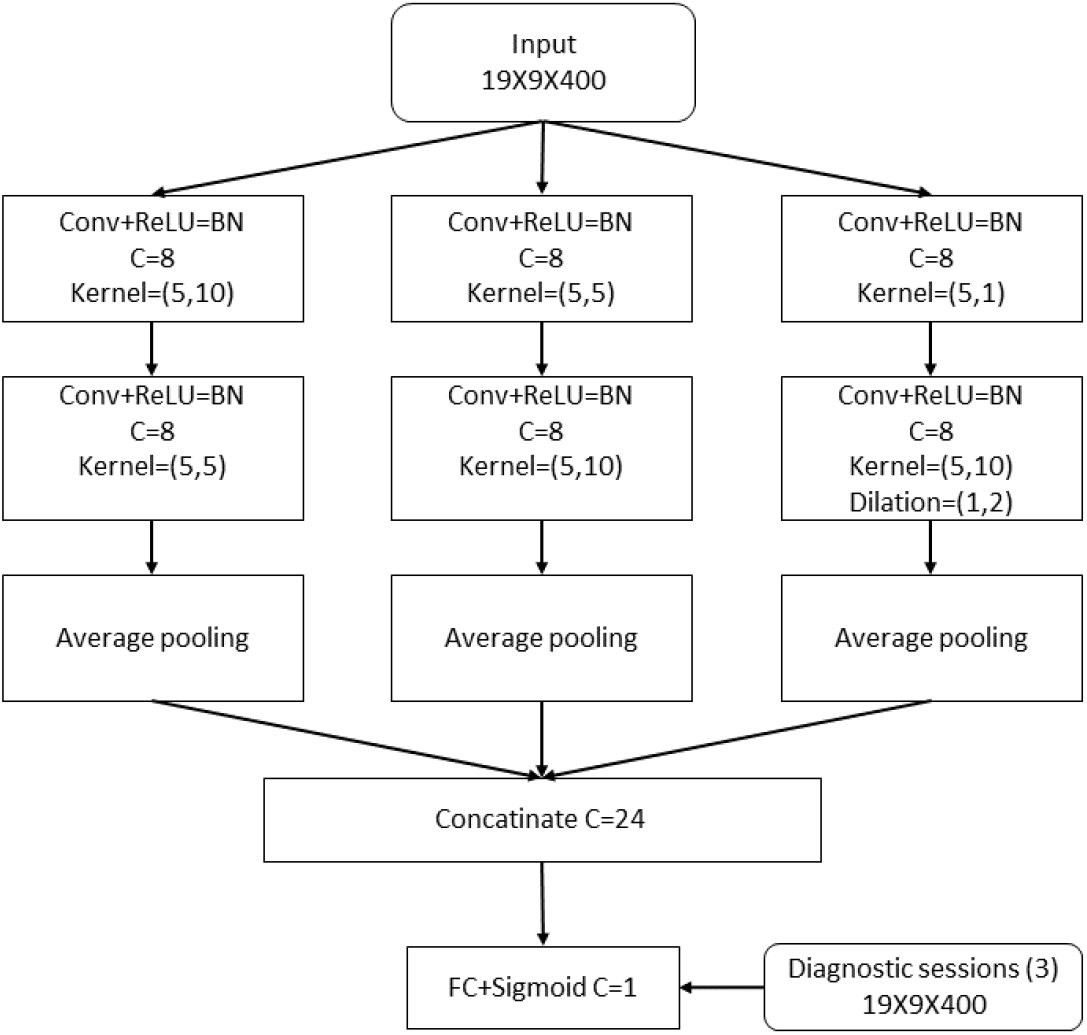
Hybrid model architecture. Operations indicated in the blocks: conv: convolution with a kernel, ReLU: rectified linear activation function, BN: batch normalization, FC: fully connected layer, C: number of channels, i.e. kernels applied at the same location. The network returns the probability that the input trial requires information retention which triggers the successful shooting down of an enemy starship.

Using time-frequency representations, post-hoc perturbation and gradient analyses showed that the EEG features driving classification of difficult (memory retention) versus easy (no memory load) trials corresponded to established neural markers of working memory.

### Pre- and post-neurofeedback training examination

Before starting and a few days after completing the diagnostic and neurofeedback training sessions all the participants performed three cognitive tests evaluating their cognitive performance mostly related to working memory. Different tasks were used than the one applied in the neurofeedback training. Each testing session started with a four-minute EEG-resting state, followed by EEG recording while the participants performed the N-back, Transitive reasoning, and Match to Sample (MTS) tasks. The scheme of all the procedures applied in the study is presented in Fig. 3

**Fig. 3.**
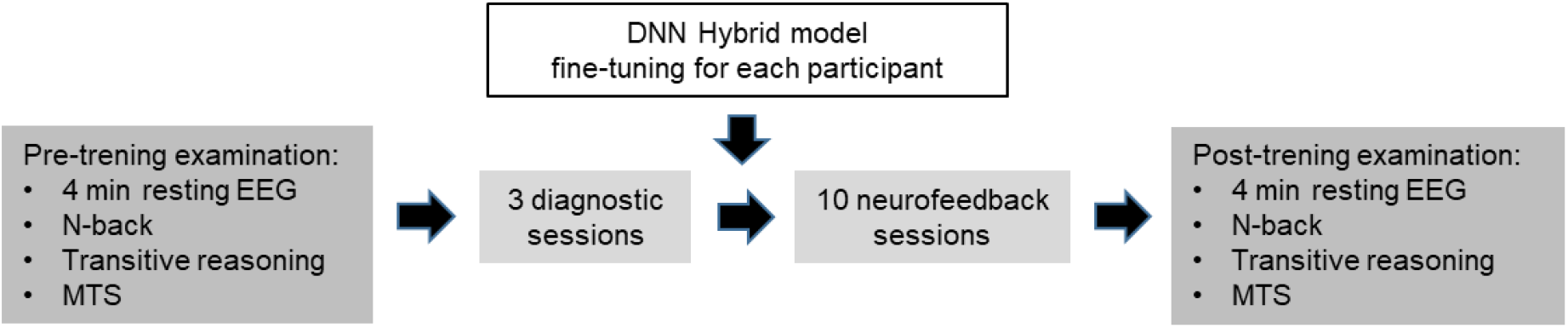
Experimental design. Scheme of the procedures performed by all participants.

### Cognitive tests

#### N-back

A visual letter variant of the N-back task [26] was used to assess working memory. The stimuli were the five letters A, B, C, D and E [27], presented in pseudorandom order. Participants were asked to react when the currently presented letter was the same as or different than the letter preceding it immediately (1-back) or either two (2-back) or three trials earlier (3-back condition). The 0-back condition required responding each time a predetermined target (X letter) appeared, and withheld responses to all other stimuli (see Fig. 4). Participants were allowed up to 2 s for each response. There were 480 stimuli (124 targets) in total. Stimulus presentation and response recording were controlled by Presentation® software (version 17.2, Neurobehavioral Systems, Inc., Berkeley, CA). Visual stimuli were displayed on a 24-inch monitor. Participants were instructed to respond quickly and accurately by pressing the left mouse button. The dependent variable comprised the d’ measure (correct hit rate minus false alarm rate). The 0-back condition was not analyzed due to strong ceiling effect.

**Fig. 4.**
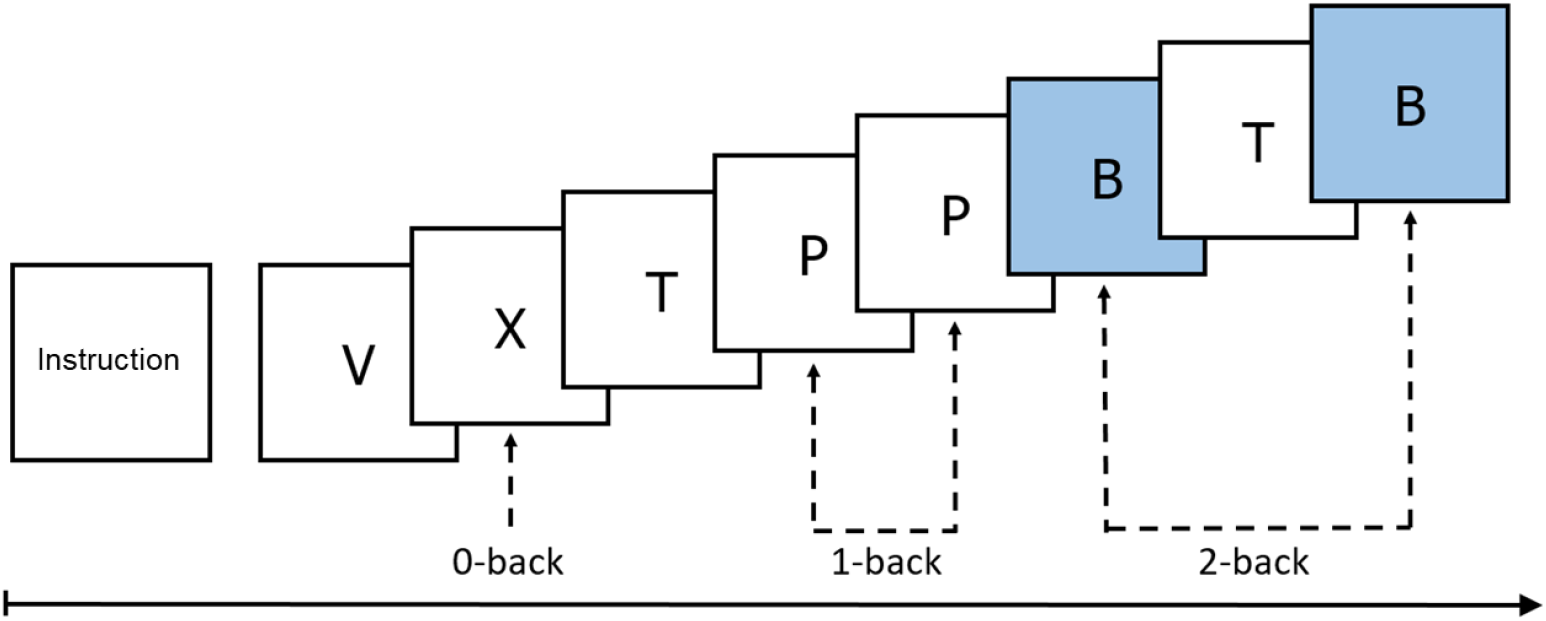
N-back task. In our procedure 0-back (X letter), 1-back, 2-back, and 3-back was performed.

#### Match to Sample task

The Match to Sample task (MTS), a simpler version of the training task, was also administered to tap into working memory. All stimuli were 4 x 4 square grids, with each square randomly colored either blue or red. For each trial, first a grid was presented in the middle of the screen for 1 s. The participants were required to memorize the arrangement of the squares in the grid. Then the grid disappeared, and after a short while two grids appeared on the screen, one on the left and the other on the right side of the screen. An example trial of the procedure is presented in Fig. 5. Participants responded with either the left or the right mouse button to indicate which of the two grids (either left or right, respectively) was the same as the grid presented previously. There were 80 trials, evenly split across two possible delay conditions (1 s and 5 s). The procedure was controlled by Presentation® software. Response accuracy was the dependent variable.

**Fig. 5.**
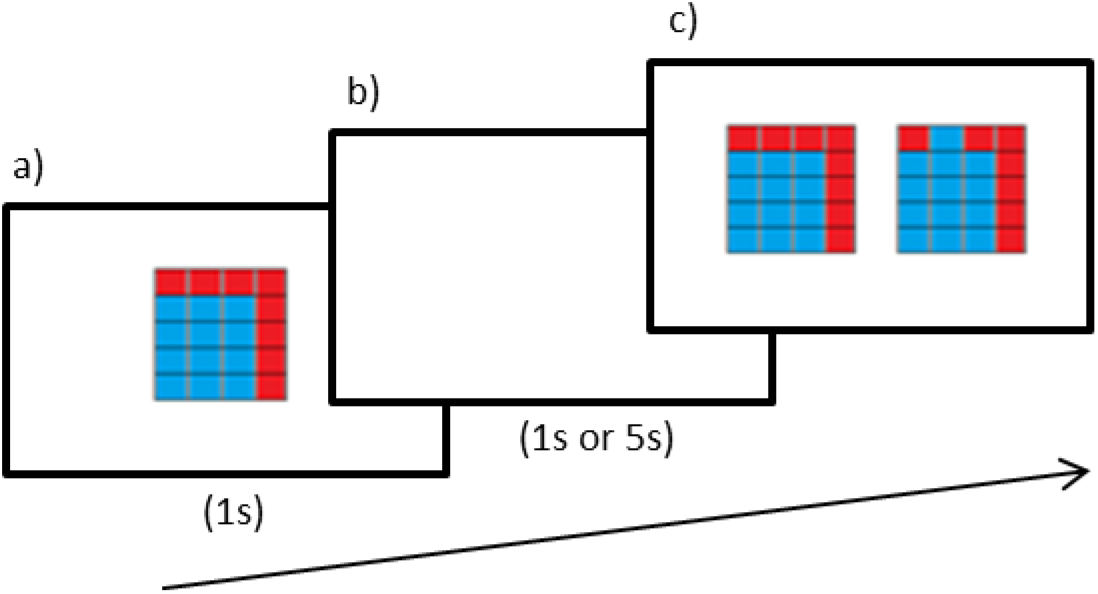
Example events in the Match-to-sample task: (a) presentation of the object to remember, (b) followed by 1 or 5 s of delay, and (c) two objects for indication which of them was presented before the delay.

#### Transitive Reasoning Task

The Transitive Reasoning Task was adapted from [28]. It aimed to tap into processing relations in working memory. The experimental procedure was developed using Presentation® software. Each trial started with the presentation of fixation cross either for 2.5 s (2/3 of trials) or for 10 s (the remaining 1/3 of trials). The latter fixation cross was aimed at recording the baseline EEG signal from the intervals interleaved with the task trials, for the purpose of some future analyses.

The task included three conditions: the easy, medium, and hard trials. In each trial, three pairs of Greek letters were presented one below another. Each pair, called a premise, was linked by either the “<” or “>” relation term (e.g., γ < φ, α > θ), meaning “smaller than” and “larger than”, respectively. The premises always reflected a univocal order of the four letters (e.g., β < λ < δ < π). In the easy trials, the premises presented the letters in the exact increasing (β < λ, λ < δ, δ < π) or decreasing (π > δ, δ > λ, λ > β) order, which thus could be relatively easily constructed out of the premises. In the medium trials, one premise was reversed: its letters were swapped, and the relation changed its direction (e.g., λ < δ reversed into δ > λ), so the order could no longer be induced from physical layout of the letters, and an additional mental operation was required. In the hard trials, two premises swapped their locations (e.g., λ < δ, β < λ, δ < π), so the entire order had to be reconstructed mentally. There were 20 trials for each condition. The order of trials and the selection of symbols were pseudo-randomized, so that no type of condition or symbols could repeat directly after each other.

First, the premises were shown for 10 s. Then, the three response options were shown below the premises (which remained at the screen), placed horizontally. Each option included two letters separated in the order by exactly one letter. They were linked by either the “<” or “>” relation term. One random option linked the letters in an order matching the order defined by the premises (e.g., β < δ), while two remaining options linked the letters in an incorrect order (e.g., β > δ, or λ > π). The task was to select the correct option that matched the order defined by the premises, by pressing an arrow button corresponding to the position of the option on the screen (←, ↓, or →). The options were shown for 10 s, or until the response was given. Seven seconds after the response options onset, a small icon of a watch was shown to inform the participants that they were running out of time. The dependent variable for each condition was the proportion of correct responses in 20 trials. Fig. 6 presents the example trial of the task.

**Fig. 6.**
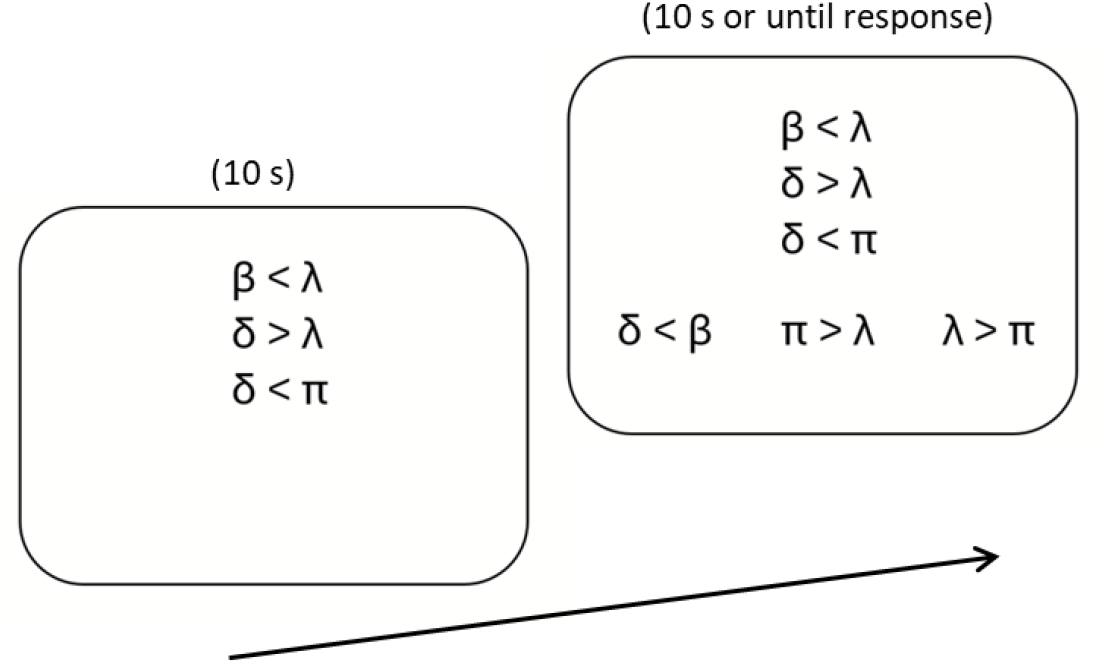
Example screen of the Transitive Reasoning Task (the medium-difficulty condition). The task was to represent mentally the increasing or decreasing order of Greek letters as defined by the three premises at the upper part of the screen, and to select the one and only correct option that matched this order, from the three response options below.

#### EEG data recording and preprocessing

In the pre- and posttest sessions, EEG data was collected during all cognitive tasks. EEG was recorded using 64 Ag/AgCl electrodes (Quick Amp; Brain Products GmbH, extended system 10-20, sampling rate 1000 Hz), referenced to FCz, and grounded at the FPz electrode. The electrode impedance was kept below 10 KΩ. The EEG data was analyzed using Matlab software (MATLAB R2020a, The MathWorks Inc, Massachusetts), EEGLAB toolbox [24], and custom Matlab scripts. Preprocessing of the EEG data included automatic artifact detection using clean_rawdata EEGLAB extension (ver. 2.3). The signal was 0.5 - 45 Hz FIR filtered and resampled to 250Hz. The *clean_rawdata* function was used to detect bad channels (correlation thresholds ranged between 0.8 to 0.5) and bad segments (amplitude threshold ranging between 10 and 100 μV) such that no more than 9 electrodes and 20% of data points were excluded.

The Transitive Reasoning Task trials were extracted based on the onset of premise stimuli, and were divided into early, mid and late phases of reasoning. Each phase was defined as a 4-second-long window with slight overlap in between: Early phases spanned 0-4 seconds from the premise onset, the mid spanned 3-7 seconds, and the late spanned 6-10 seconds. After trial extraction, Infomax ICA was performed. Using IClabel EEGLAB’s extension (Pion-Tonachini et al., 2019), automatic classification and removal of bad components was applied. The thresholds for removal were set to 70% for the eye, 80% for the ECG, 80% for the muscle, and 80% for the bad channel type of artifacts. After the removal of ICs, the deleted channels were interpolated, re-referenced to signal averaged over all electrodes, and baseline corrected (by removal of the mean of all channels in each trial). The data was then visually inspected for any remaining bad trials. For consistency, resting-state EEG data analysis followed an identical preprocessing pipeline, except that the dummy markers were inserted every 4 seconds instead of trial extraction based on actual events during the task.

#### EEG data analysis

The neurofeedback protocols (i.e., EEG parameters such as electrodes, frequencies, powers and signal correlations) used in the experiment resulted from personalized fine tuning of the pre-trained artificial neuronal networks [18]. The parameters of each participant’s neurofeedback protocol were uniquely encoded within the DNN overseeing the training process and were not accessible to participants, trainers, or experimenters. Consequently, our analysis was limited to evaluating whether EEG activity changes linked to correct responses exhibited any consistent patterns across individuals. We hypothesized that effective training would result in the reorganization of brain networks involved in working memory processes underlying task performance. To explore this, we assessed changes in brain network organization through graph-theoretical measures and power spectrum analyses.

To this end we analysed the power and global graph metrics in the power spectrum for consecutive EEG bands (Delta: 1.0–3.5 Hz, Theta: 4.0–7.0 Hz, Alpha: 8.0–13.0 Hz, Low-Beta: 14.0–20.0 Hz, High-Beta: 20.0–26.0 Hz and Low-Gamma: 30.0–45.0 Hz). Data from one participant was excluded from further analysis due to poor performance. The final set included data from 28 participants who completed the training procedure and 28 participants from the control group.

#### Power spectrum analyses

Power was calculated by computing the Power Spectral Density (PSD) using the Welch method [30]. Specifically, PSD was computed for both pre- and posttest sessions in the frequency range of interest. For each session, the PSD was averaged across the selected electrodes. The results were expressed in microvolt squared (µV^2^). The power analyses aimed at testing whether the training intervention affected the power at any band in the experimental group, as compared to the control group. Electrodes were grouped into four clusters **Frontal**: Fp1, Fp2, AF7, AF3, AFz, AF4, AF8, F7, F5, F3, F1, Fz, F2, F4, F6, F8, **Parietal**: P7, P5, P3, P1, Pz, P2, P4, P6, P8, PO7, PO3, POz, PO4, PO8, **Left temporal**: FT7, FC5, T7, C5, TP7, CP5 and **Right temporal**: FT8, FC6, T8, C6, TP8, CP6. The electrode clusters were selected following commonly used groupings in EEG research to provide representative coverage of frontal, parietal, and temporal regions of interest (e.g.,[31]). In addition, electrode clusters were defined to ensure sufficient spatial sampling within each region, while reducing the dimensionality of statistical analyses.

#### Graph metrics analysis

The brain’s structural and functional systems have features of complex networks [32], which could be estimated using the graph theory, providing metrics necessary to characterize complex networks [33], using functional connectivity matrix [34]. Consequently, we aimed at assessing how the reinforcement of the EEG activity associated with correct responses was translated into the brain network organization. We used two basic measures: modularity, which is a statistic that quantifies the degree to which the network may be subdivided into clearly delineated groups, and global efficiency, which quantifies the average minimal distance of a node to all other nodes in the network. Metrics were calculated using the Brain Connectivity Toolbox, originally developed by Rubinov and colleagues ([34],[35]), implemented in Python (LaPlante, 2023). For calculation of the functional connectivity matrix we used weighted Phase Lag Index (wPLI), an extension of the Phase Lag Index [36], which quantifies the consistency of phase lead-lag relationships between signals from different brain regions. Unlike simple coherence, wPLI reduces the unwanted influence of volume conduction and common sources, by focusing only on the non-zero phase lag components. It provides insights into true functional connectivity, reflecting synchronized neural interactions that likely represent genuine communication patterns between brain areas, which makes it a suitable tool for graph analysis [37]. The wPLI values range 0 to 1, where 0 indicates no consistent phase-lagged interaction between the two signals, suggesting either no functional connectivity or the spurious one that likely reflects noise or volume conduction, and 1 signifies perfectly consistent phase-lagged interaction, meaning that the phase difference between signals is maximally stable across time. The latter indicates strong, directionally consistent neural coupling. Analyses were performed individually for each task trial type and graph metrics. Analyses were conducted for 28 participants from the experimental group and 27 participants from the control group, as one participant had to be excluded from the latter group due to unreliable EEG data.

#### Statistical analyses

All the analysis of the EEG data was conducted using Python [38] and the MNE library [39]. For power spectrum analyses each band and each electrode cluster were tested using a two-way interaction of Group (training vs. control) and Session (pretest vs. posttest), expecting that the power change between the pretest and the posttest would be significantly larger in the training than in the control group. For graph metrics, the Shapiro-Wilk test was first applied to assess normality. If the data followed a normal distribution, a two-way mixed ANOVA with post-hoc tests (implemented in the Pingouin Python library) was conducted. Otherwise, Mann-Whithney two-sided pairwise test was used. In the transitive reasoning task, EEG data from all three trial variants (easy, medium, and hard) were pooled to enhance statistical power.

## Results

### Behavioral scores on working memory tasks

First, we checked whether active training affected posttest performance on the two working memory tasks: the MTS and N-back, relative to the sham control group. In the former task, there was a significant main effect of the Session, *F*(1, 55) = 4.25, *p* = .044, η^2^ = .07, suggesting improved accuracy from the pretest (M ± SE: 77.2% ± 1.5%) to posttest (80.1% ± 1.2%). However, the interaction with the Group factor was not significant, *F*(1, 55) = 0.019, p = .891, indicating that the Active Training and the Sham Control Group improved accuracy comparably.

In the N-back task, a significant effect of Session (pretest *d*’ ± SE: 73.6% + 1.0%, posttest *d*’ = 75.7% + 1.1%), *F*(1, 54) = 4.89, *p* = .031, η^2^ = .08, did not translate into a significant two-way interaction with the Group factor either, *F*(1, 54) = 1.34, *p* = .253. For none of the N conditions, there was a significant Group-Session contrast, each *p* > .11.

Accuracy on the Transitive Reasoning Task was submitted to 2×2×3 ANOVA (Group: training vs. control; Session: pretest vs. posttest; Condition: easy, medium, hard). Each factor yielded a statistically significant effect: The training group displayed higher accuracy (M ± SE: 53.6% ± 3.5%) than the control group (43.6% ± 3.5%), *F*(1, 54) = 4.06, *p* = .048, η^2^ = .07, accuracy was higher at the posttest (52.0% ± 2.6%) than at the pretest (45.3% ± 2.5%), *F*(1, 54) = 25.28, p < .001, η^2^ = .37, and it decreased from the easy (54.1% ± 2.9%), through the medium (47.0% ± 2.6%), to the hard trials (44.9% ± 2.5%), *F*(2, 108) = 14.28, p = .002, η^2^ = .21.

These main effects were qualified by two significant two-way interactions. First, the increase in accuracy between the pretest and posttest was larger for the easy items (Δ11.9% ± 3.1%) than for the medium (Δ5.4% ± 2.8%) and the hard items (Δ2.7% ± 2.7%), *F*(2, 108) = 6.48, *p* = .002, η^2^ = .11. Crucially, a significant interaction of Group and Session, *F*(1, 54) = 8.37, *p* = .006, η^2^ = .13, indicated that the training group increased accuracy between the pretest and posttest to a larger extent (Δ12.6% ± 4.6%) than did the control group (Δ2.9% ± 3.6%). Planned comparisons showed that the former gain was highly significant, *F*(1, 54) = 31.36, *p* < .001, whereas the latter gain was not, *F*(1, 54) = 2.28, *p* = .138. Although the three-way interaction was not significant, *F*(2, 108) = 1.49, *p* = .229, η^2^ = .03, analysis of contrasts even further differentiated the gains in the training vs. control group (see Fig.7). Specifically, the latter group significantly increased their accuracy from the pretest to posttest only in the easy trials, *p* < .001, but not in the two remaining conditions, each *p* > .40. By contrast, the gains in the training group were highly significant in all three conditions, each *p* < .01. Moreover, at the pretest, the groups did not differ in the easy, medium, and hard trials, *p* = .07, *p* = .44, and *p* = .56, respectively, while at the posttest the training group surpassed the control group in all the three conditions, each *p* < .03.

**Fig. 7.**
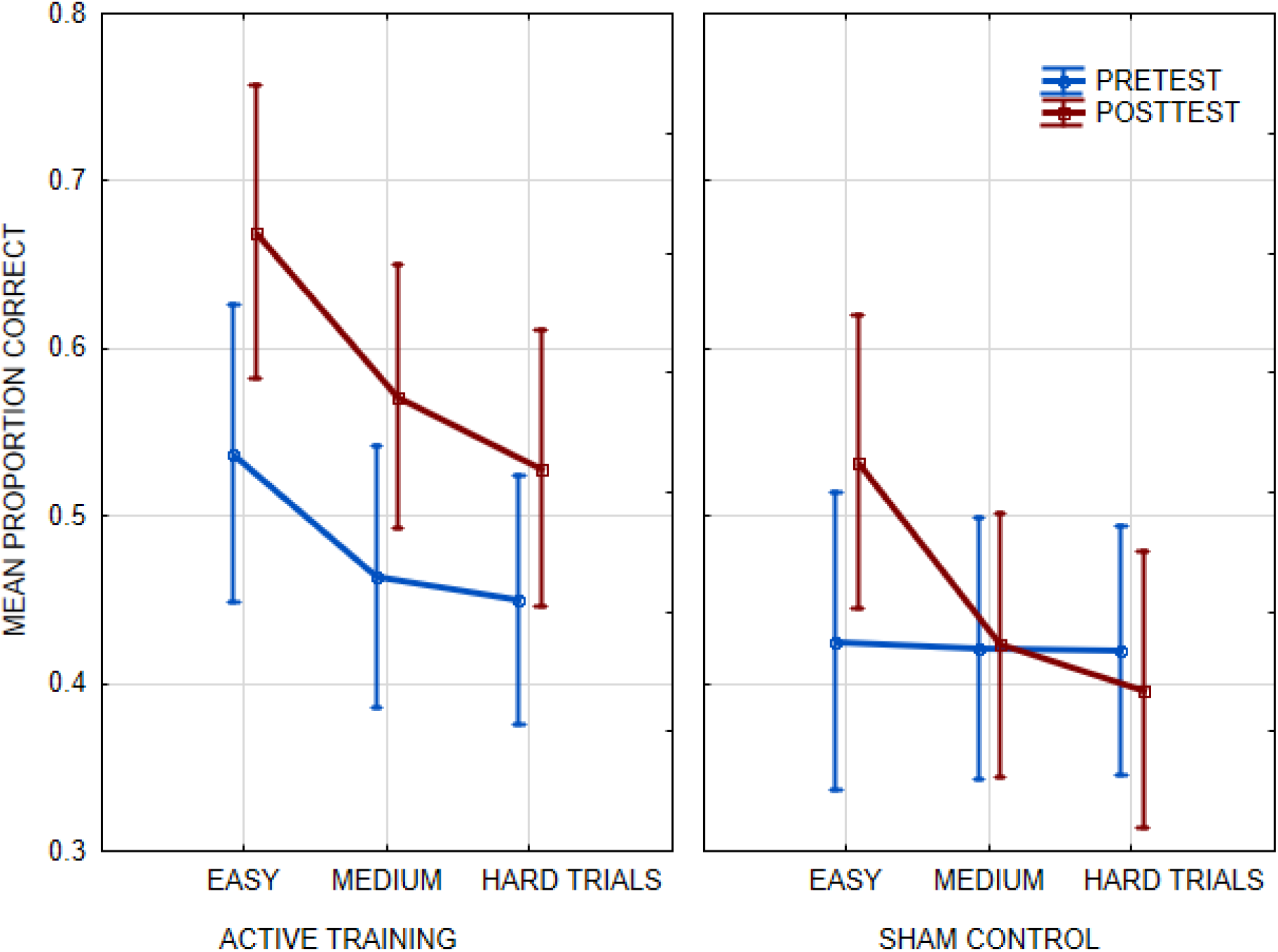
Mean accuracy in the transitive reasoning task shown separately for the pretest and posttest in the active training and the sham control group, for the easy, medium, and hard trials. Vertical bars indicate 95% confidence intervals.

### EEG analyses

#### Pre- and post-training eyes open resting state analyses

None of the comparisons showed significant differences prior to or post the neurofeedback intervention, including power spectral density across major EEG frequency bands, binary wPLI connectivity comparisons, and connectivity based graph-theoretical metrics.

#### Task related EEG data analyses

Based on the behavioral outcomes of the administered tasks, we chose to analyze only the data from tasks that demonstrated statistically significant group effects. As a result, EEG data analysis was limited to the Transitive Reasoning Task.

##### Alpha Band Power

An interaction between Group and Session was observed for the alpha band power, *F*(1, 53) = 6.39, *p* = 0.014, η^2^ = .11. This effect was driven by changes in the parietal cluster, with no significant differences found in the frontal, *p* = .103, left temporal, *p* = 145, and right temporal cluster, *p* = .169. Specifically, the control group showed significant increase in the parietal alpha power from the pretest to posttest, *F*(1, 53) = 5.62, *p* = 0.021, whereas the alpha power in the training group did not significantly change, *F*(1, 53) = 1.43, *p* = 0.238. These findings are shown in Fig. 8. There was also a non-significant trend towards the Group × Session interaction for the theta band in the left temporal cluster, *F*(1, 53) = 3.45, *p* = 0.069, η^2^ = .06. No other frequency band change from the pretest to posttest significantly differentiated the two groups, each *p* > .17.

**Fig. 8.**
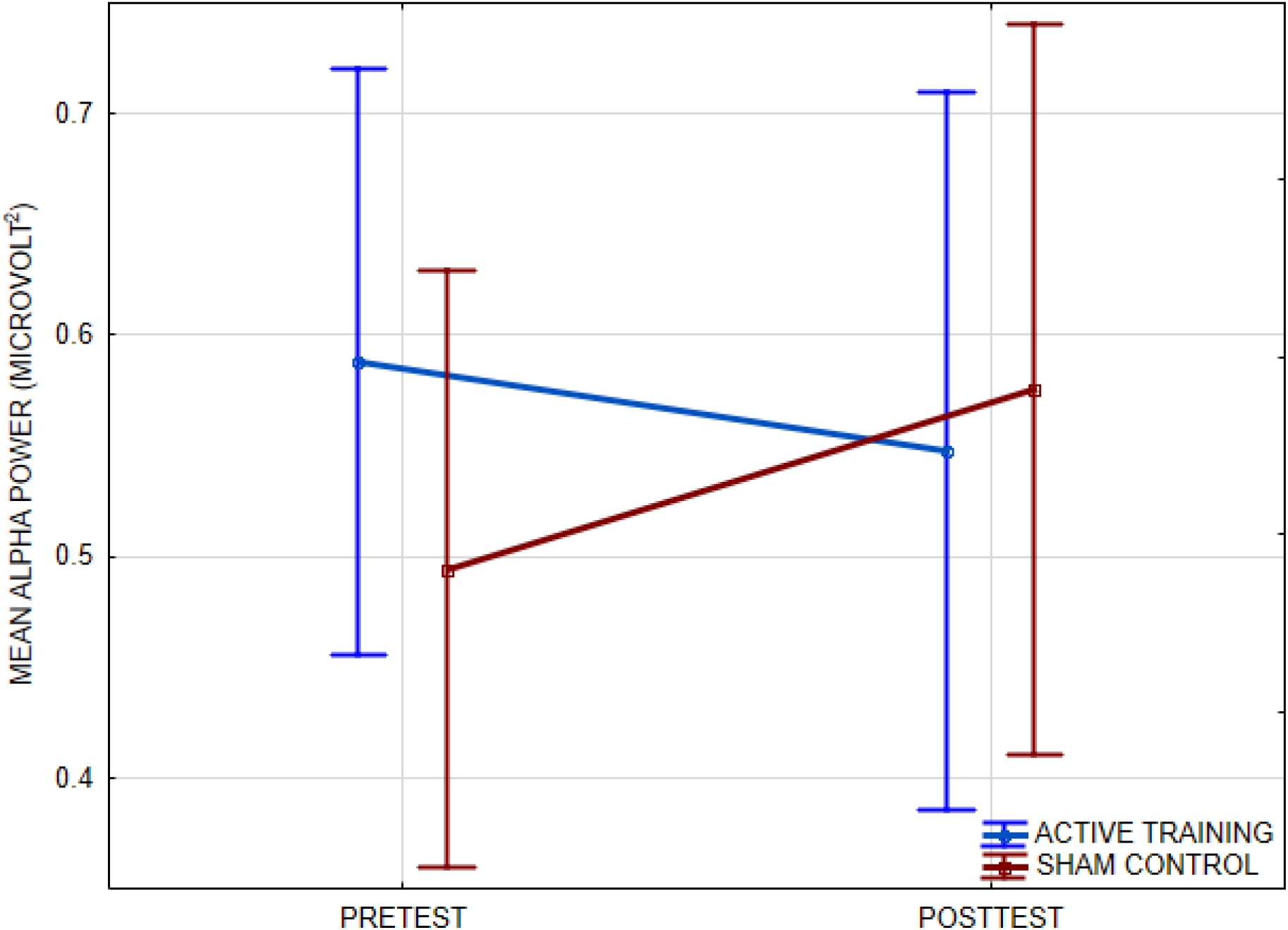
Mean alpha power in EEG data recorded in the Transitive Reasoning Task shown separately for the pretest and the posttest in the active training and the control sham group. Vertical bars indicate 95% confidence intervals.

#### Overall band-wise estimation of wPLI

Finally, we compared average wPLI values between the training and the control group for the six above defined bands. To this aim, we calculated wPLI for 11 pairs of neighbouring electrodes within the medial frontal and parietal cluster of electrodes (the following electrode pairs: Fp1-Fp2, Fp1-F3, Fp2-F4, F3-Fz, F4-Fz, Fz-Pz, F3-P3, F4-P4, Pz-P3, Pz-P4, P3-P4), for the pretest and posttest Transitive Reasoning Task data. This cluster is widely recognized as most strongly associated with reasoning and cognitive ability (Haier, 2017). Next, the corresponding pretest wPLI values were subtracted from the consecutive posttest Transitive Reasoning Task wPLI values, with the outcome value reflecting the change in the wPLI values from pretest to posttest. Finally, the 11 wPLI values for one and the same band were averaged, yielding 6 estimates (one per band) of the overall information transfer change related to performance on the Transitive Reasoning Task. These values are shown in Fig. 9, separately for the training and the control group. There was a significant two-way interaction between the Group and the Band factors, *F*(5, 265) = 2.79, *p* = .017, η^2^ = 0.050, resulting from a larger change of wPLI in the training group, relative to the control group, in the low- and high beta band as well as the low-gamma band, while the corresponding wPLI values in the delta, theta, and alpha band were comparable between the two groups. Specifically, contrasts indicated the training group’s significantly larger change in the high-beta band, *F*(1, 53) = 5.93, *p* = .018, and an analogous trend in the low-beta, *F*(1, 53) = 3.29, *p* = .075, and low-gamma band, *F*(1, 53) = 3.47, *p* = .068, but no difference in the delta, theta, and gamma bands, *p* = .845, *p* = .549, and *p* = .455, respectively. These results suggested that any effects of training in connectivity patterns were most likely to be found in the beta and gamma frequency bands.

**Fig. 9.**
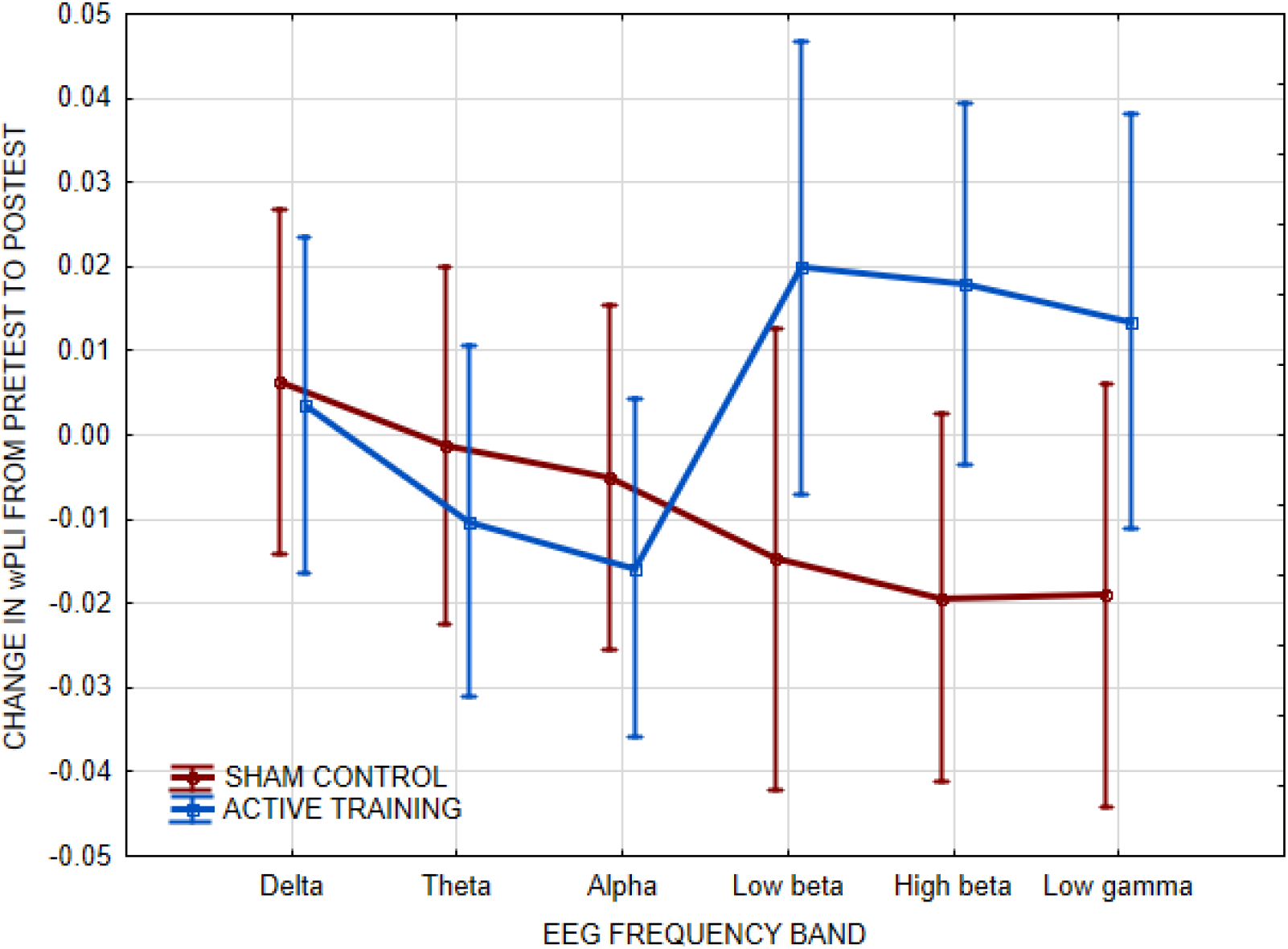
A change in mean connectivity (weighted Phase Lag Index, wPLI) in the EEG data between the pretest and posttest of the Transitive Reasoning Task in the medial frontal and parietal electrode cluster, for the active training vs. sham control group.

## Discussion

In summary, our Task-Pretrained, Subject-Finetuned Neurofeedback (TPSF-NF) framework was tested on adults between 41 and 64 years old in order to enhance their working memory. We found that the training group showed significantly greater improvement from the pretest to posttest on the Transitive Reasoning Task than the control group. In contrast, we observed no significant group differences on the two remaining working memory tasks (the N-back and Match-to-Sample).

Given the absence of transfer to these two working memory tasks, we next examined EEG dynamics during the Transitive Reasoning Task to test whether behavioral gains were accompanied by neural changes. We analyzed two canonical EEG markers: band power and wPLI. Despite individually tailored protocols, we observed a consistent pattern: the training group showed lower alpha-band power (~10 Hz), reflecting greater alpha desynchronization compared to controls. Greater alpha desynchronization is typically interpreted as a shift toward increased processing in other frequency ranges (theta, beta, and possibly gamma; [40], [41]). In line with this finding, we observed increased beta-band (~25 Hz) phase coherence, reflecting more focused and coordinated neural processing ([42], [36]). Therefore, we can hypothesise that our TPSF-NF protocol presumably led to the enhancement of neural control and coordination of relatively faster brain rhythms that persisted after the training and resulted in improved cognitive processing on a reasoning task.

Individually tailored protocols may produce heterogeneous effects across tasks and patients. Such variability is not surprising, given that working memory performance can be influenced by multiple factors (e.g., neural pathway integrity, motor function). Nevertheless, the absence of transfer to the two relatively simple working memory tasks, under the robust effect for a more complex task, is surprising. It is difficult to speculate on potential reasons underpinning this null outcome, but two hypotheses can be considered. First, working memory tasks seem to rely on the neural activity at the beta band to a smaller extent than does reasoning, as the leading neurophysical models of working memory ([43]), as well as their empirical verification ([44], [45], [46]), all point at the theta (~5 Hz) and gamma oscillations (>40 Hz) as crucial for working memory maintenance. It is possible that our TPSF-NF training protocol did not target these two bands, as suggested by the results for the band power and wPLI. By contrast, several studies suggested a role of the 20-30 Hz frequency band in reasoning ([47], [31]).

Second, even though our protocol targeted neural processing overlapping to some extent with working memory, the two working memory tasks we applied might have been too simple to capture such neural enhancement. Specifically, both tasks required recognition of stimulus occurrence (the same/different), but not precise recollection of its identity and attributes. The former process can often rely solely on the feeling of stimulus familiarity, which can be fairly automatic and may not involve working memory mechanisms to a large extent ([48], [49]), unlike tasks which would demand full recollection [50].

Our review of recent neurofeedback studies shows that only a limited number of such studies have applied machine learning methods. Most of them focused either on choosing suitable models for neurofeedback [51] or on applications outside cognitive performance, such as pain management [52,53]. The study most comparable to ours is [54]. They used a Common Spatial Pattern (CSP) method for individual feature extraction and Linear Discriminant Analysis (LDA) for supervised training. However, they reported the experimental group’s poorer recognition accuracy in the posttest, while no changes were observed in the control group.

Several methodological differences may explain these divergent results. First, their design relied on a single diagnostic and training session, whereas ours spanned multiple sessions across different days. Second, they used CSP and LDA, while we employed DNNs. Finally, they evaluated outcomes on a task highly similar to their training task, while we assessed transfer to a reasoning task. These distinctions, alone or combined, likely contributed to the improvements observed in our study.

In conclusion, our findings demonstrate that the neurofeedback training protocol based on task-pretrained DNNs, further fine-tuned to individuals, can lead to improvements in cognitive performance which span beyond the trained task, as well as to corresponding increases in neural efficiency. This highly original result (i.e., no neurofeedback so far has succeeded with respect to complex cognition (see [54]) yields important ramifications for the development of effective cognitive enhancement protocols, especially those aimed at middle-aged and older adults, which may have potential to prevent cognitive decline and preserve cognitive reserve [55]. Future studies should validate the effectiveness of our neurofeedback framework also for broader domains of cognitive enhancement beyond working memory/reasoning.

### Limitations and future directions

This study has several limitations. The most important one pertains to the absence of follow-up assessments after delay periods, which would clarify for how long the training effects persisted and whether booster sessions would be needed or not. Regarding the criterion tasks, more complex and difficult working memory tasks, as compared to the N-back and Match-to-Sample, should have been also administered (e.g., complex span tasks; see [50]. Another limitation concerns the lack of fine-grained analysis of individualized training protocols. While our neural network framework supports interpretability through feature importance analysis, a full examination of individual training trajectories is beyond this study and requires a dedicated investigation. Nevertheless, the current study likely comprises a good starting point for an application of individualized machine learning-based neurofeedback training in enhancement of various cognitive and brain functions, with a potentially high impact on the quality of life in selected populations.

## Notes

### Competing Interest Statement

The authors have declared no competing interest.

### Funding

- Mazovian Regional Financing UnitRegional Financing Unit: RPMA.01.02.00-14-b459/18-00

## References

[1] V.L. Feigin, et. al, Lancet Neurol. 16 (2017) 877–897.

[2] N. Bittner, C. Jockwitz, T.W. Mühleisen, F. Hoffstaedter, S.B. Eickhoff, S. Moebus, U.J. Bayen, S. Cichon, K. Zilles, K. Amunts, S. Caspers, Nat. Commun. 10 (2019) 621.

[3] S.M. Courtney, T. Hinault, Prog. Neurobiol. 203 (2021) 102076.

[4] J.I. Fleck, M. Arnold, B. Dykstra, K. Casario, E. Douglas, O. Morris, Front. Aging Neurosci. Volume 11-2019 (2019).

[5] D.V.P.S. Murty, K. Manikandan, W.S. Kumar, R.G. Ramesh, S. Purokayastha, M. Javali, N.P. Rao, S. Ray, Neuroimage 215 (2020) 116826.

[6] C. Babiloni, G. Binetti, A. Cassarino, G. Dal Forno, C. Del Percio, F. Ferreri, R. Ferri, G. Frisoni, S. Galderisi, K. Hirata, B. Lanuzza, C. Miniussi, A. Mucci, F. Nobili, G. Rodriguez, G. Luca Romani, P.M. Rossini, Hum. Brain Mapp. 27 (2006) 162–172.

[7] E.Y. Privodnova, N. V Volf, Hum. Physiol. 42 (2016) 469–475.

[8] F.H. Duffy, G.B. McAnulty, M.S. Albert, Neurobiol. Aging 14 (1993) 73–84.

[9] M. Thomas, Int. J. Psychophysiol. 101 (2016) 33–42.

[10] Fabrizio Vecchio, Francesca Miraglia, Placido Bramanti, Paolo Maria Rossini, J. Alzheimer’s Dis. 41 (2014) 1239–1249.

[11] R. Chow, R. Rabi, S. Paracha, L. Hasher, N.D. Anderson, C. Alain, Neuroscience 485 (2022) 116–128.

[12] D.F. Hultsch, C. Hertzog, B.J. Small, L. McDonald-Miszczak, R.A. Dixon, Psychol. Aging 7 (1992) 571–584.

[13] D.C. Park, G. Lautenschlager, T. Hedden, N.S. Davidson, A.D. Smith, P.K. Smith, Psychol. Aging 17 (2002) 299–320.

[14] L. Nyberg, M. Lövdén, K. Riklund, U. Lindenberger, L. Bäckman, Trends Cogn. Sci. 16 (2012) 292–305.

[15] M. Garcia Pimenta, T. Brown, M. Arns, S. Enriquez-Geppert, Neuropsychiatr. Dis. Treat. 17 (2021) 637–648.

[16] R.T. Schirrmeister, J.T. Springenberg, L.D.J. Fiederer, M. Glasstetter, K. Eggensperger, M. Tangermann, F. Hutter, W. Burgard, T. Ball, Hum. Brain Mapp. 38 (2017) 5391–5420.

[17] F. Lotte, L. Bougrain, A. Cichocki, M. Clerc, M. Congedo, A. Rakotomamonjy, F. Yger, J. Neural Eng. 15 (2018) 031005.

[18] J. Żygierewicz, R.A. Janik, I.T. Podolak, A. Drozd, U. Malinowska, M. Poziomska, J. Wojciechowski, P. Ogniewski, P. Niedbalski, I. Terczynska, J. Rogala, J. Neural Eng. 19 (2022) 046053.

[19] F. Faul, E. Erdfelder, A.-G. Lang, A. Buchner, Behav. Res. Methods 39 (2007) 175–191.

[20] Y.-H. Tseng, K. Tamura, T. Okamoto, Sci Rep 11 (2021) 17274.

[21] R. Rozengurt, I. Kuznietsov, T. Kachynska, N. Kozachuk, O. Abramchuk, O. Zhuravlov, A. Mendelsohn, D.A. Levy, Cogn. Affect. Behav. Neurosci. 23 (2023) 1473–1481.

[22] W. Zhou, W. Nan, K. Xiong, Y. Ku, Npj Sci. Learn. 9 (2024) 32.

[23] L. Shen, Y. Jiang, F. Wan, Y. Ku, W. Nan, Neurobiol. Learn. Mem. 205 (2023) 107834.

[24] U. Malinowska, J. Wojciechowski, M. Waligóra, J. Rogala, Front. Physiol. 15 (2024) 1457371.

[25] S. Ioffe, C. Szegedy, F. Bach, D. Blei, in: PMLR (Ed.), Proc. 32nd Int. Conf. Mach. Learn., Lille, France, 2015, pp. 448–456.

[26] A. Gevins, B. Cutillo, Electroencephalogr. Clin. Neurophysiol. 87 (1993) 128–143.

[27] T.E. Kearney-Ramos, J.S. Fausett, J.L. Gess, A. Reno, J. Peraza, C.D. Kilts, G.A. James, J. Int. Neuropsychol. Soc. 20 (2014) 736–750.

[28] A. Chuderski, Intelligence 77 (2019) 101396.

[29] A. Delorme, S. Makeig, J. Neurosci. Methods 134 (2004) 9–21.

[30] P. Welch, IEEE Trans. Audio Electroacoust. 15 (1967) 70–73.

[31] M. Ociepka, P. Kałamała, A. Chuderski, Intelligence 100 (2023) 101780.

[32] D.S. Bassett, E.T. Bullmore, Curr. Opin. Neurol. 22 (2009).

[33] E. Bullmore, O. Sporns, Nat. Rev. Neurosci. 10 (2009) 186–198.

[34] M. Rubinov, O. Sporns, Neuroimage 52 (2010) 1059–1069.

[35] M. Rubinov, R. Kötter, P. Hagmann, O. Sporns, Neuroimage 47 (2009) S169.

[36] C.J. Stam, G. Nolte, A. Daffertshofer, Hum. Brain Mapp. 28 (2007) 1178–1193.

[37] M. Hardmeier, F. Hatz, H. Bousleiman, C. Schindler, C.J. Stam, P. Fuhr, PLoS One 9 (2014) e108648.

[38] G. Van Rossum, F.L. Drake, Python 3 Reference Manual: (Python Documentation Manual Part 2), CreateSpace Independent Publishing Platform, 2009.

[39] A. Gramfort, M. Luessi, E. Larson, D.A. Engemann, D. Strohmeier, C. Brodbeck, R. Goj, M. Jas, T. Brooks, L. Parkkonen, M. Hämäläinen, Front. Neurosci. Volume 7-2013 (2013).

[40] W. Klimesch, Brain Res. Rev. 29 (1999) 169–195.

[41] G. Pfurtscheller, A. Aranibar, Electroencephalogr. Clin. Neurophysiol. 46 (1979) 138–146.

[42] A. V Sazonov, C.K. Ho, J.W.M. Bergmans, J.B.A.M. Arends, P.A.M. Griep, E.A. Verbitskiy, P.J.M. Cluitmans, P.A.J.M. Boon, Biol. Cybern. 100 (2009) 129–146.

[43] J.E. Lisman, M.A. Idiart, Science 267 (1995) 1512–5.

[44] A. Bahramisharif, O. Jensen, J. Jacobs, J. Lisman, PLoS Biol. 16 (2018) e2003805.

[45] M. Leszczyński, J. Fell, N. Axmacher, Cell Rep. 13 (2015) 1272–1282.

[46] S.E. Qasim, I. Fried, J. Jacobs, Cell 184 (2021) 3242-3255.e10.

[47] A. Gągol, M. Magnuski, B. Kroczek, P. Kałamała, M. Ociepka, E. Santarnecchi, A. Chuderski, Intelligence 66 (2018) 54–63.

[48] L.L. Jacoby, J. Mem. Lang. 30 (1991) 513–541.

[49] A.P. Yonelinas, M. Aly, W.-C. Wang, J.D. Koen, Hippocampus 20 (2010) 1178–1194.

[50] N. Unsworth, R.W. Engle, Psychol. Rev. 114 (2007) 104–132.

[51] A. Mockevičius, A. Voicikas, V. Jurkuvėnas, P. Tarailis, I. Griškova-Bulanova, Appl. Psychophysiol. Biofeedback (2024).

[52] M.G.M. Saif, L. Sushkova, M. Fraser, SN Comput. Sci. 4 (2023) 598.

[53] M.G.M. Saif, M.A. Hasan, A. Vuckovic, M. Fraser, S.A. Qazi, SN Appl. Sci. 3 (2021) 58.

[54] G. Campos-Arteaga, J. Flores-Torres, F. Rojas-Thomas, R. Morales-Torres, D. Poyser, R. Sitaram, E. Rodríguez, S. Ruiz, Int. J. Psychophysiol. 203 (2024) 112406.

[55] C. Pettigrew, A. Soldan, Curr. Neurol. Neurosci. Rep. 19 (2019) 1.

